# Caspase-dependent ablation of indirect medium spiny neurons projecting to external globus pallidus promotes compulsive ethanol-seeking and drinking behaviors

**DOI:** 10.1101/2025.02.21.639573

**Authors:** Humza Haroon, Matthew Baker, Doo-Sup Choi

## Abstract

The dorsomedial striatum (DMS) is primarily recognized for regulating goal-directed reward-seeking behaviors, while the dorsolateral striatum (DLS) is predominantly associated with movement and habitual behaviors. In this study, we sought to investigate two pathways, direct medium spiny neuron (dMSN) and indirect medium spiny neuron (iMSN) in the two dorsal striatal subregions (DMS and DLS) in ethanol-seeking and drinking behaviors. Here, we selectively ablated iMSN^DMS→GPe^ and iMSN^DLS→GPe^ and trained mice to exhibit goal-directed and habitual reward-seeking behaviors using random ratio (RR) and random interval (RI) operant conditioning, respectively. We found that partial ablation of iMSN^DLS→GPe^ exhibited increased resilience to bitter-tasting ethanol solution when subjected to quinine adulterated reward in operant conditioning paradigms, suggesting compulsive-like seeking behavior. Consistently, in a separate cohort of mice, we found that the iMSN^DLS→GPe^ ablated mice show higher preference and consumption of quinine adulterated ethanol solution than control mice in two-bottle choice continuous access drinking, with increasing quinine concentration and exhibit more compulsive-like behavior. On the other hand, ablation of iMSN^DMS→GPe^ resulted in insensitivity to satiety-based reward devaluation in RR-trained mice, consistent with a shift toward habitual behavior but no change in compulsive ethanol-seeking and drinking behavior. Together, our findings demonstrate that DLS iMSN function is essential in inhibiting compulsive-like behavior.

## Introduction

The basal ganglia are a group of subcortical nuclei, of which coordinated activity allows decision-making, reward-related learning and motor actions (Schultz et al., 1997; Packard and Knowlton, 2002; Yin and Knowlton, 2006). In humans, the dorsal striatum (DS) is anatomically separated by the internal capsule forming the caudate and putamen nucleus whereas, in rodents, DS is a homogeneous structure with overlapping axonal connectivity due to an absence of anatomical border (Hoover et al., 2003; Alloway et al., 2006; Alloway et al., 2009; Hooks et al., 2018). However, the rodent DS has two functionally recognizable subregions, the dorsomedial striatum (DMS, equivalent to caudate in primates and humans) and the dorsolateral striatum (DLS, equivalent to putamen) (Hauber and Schmidt, 1994; Graybiel, 2008). The DMS plays an essential role in the acquisition and expression of goal-directed behavior controlled by action-outcome (A-O) contingency, while the DLS is primarily involved in habitual behaviors regulated by stimulus-response (S-R) association (Voorn et al., 2004; Yin and Knowlton, 2006; Graybiel, 2008).

The striatum consists of GABAergic medium spiny neurons (MSN), making up more than 95% of the neuronal population, divided into equal proportions of direct pathway MSN (dMSN) and indirect pathway MSN (iMSN) (Gerfen and Surmeier, 2011). These two morphologically indistinguishable striatal MSN have extensive dendritic arbors providing synaptic integration from upstream neurons (Kawaguchi et al., 1990). In the classic model of basal ganglia, direct and indirect pathways have been considered to play opposing roles, with the direct pathway sending the “go signal” for action promotion, while the indirect pathway sends the “no go signal” to inhibit or suppress the action (Albin et al., 1989; Alexander and Crutcher, 1990; DeLong, 1990; Mink, 2018). In general, dMSN stimulation enhances reinforcement learning, whereas iMSN stimulation induces avoidance behaviors and inhibits reinforcement (Lobo and Nestler, 2011; Kravitz et al., 2012). However, accumulating evidence suggests that both dMSN and iMSN are co-activated during movement and action selection (Cui, 2013; Tecuapetla et al., 2014). This suggests that arbitration of the activities between the two MSNs determines the direction of ongoing actions. Consequently, a higher iMSN activation is associated with action termination (Cui, 2013; Geddes et al., 2018). Although studies have investigated the distinct DMS vs. DLS or iMSN vs. dMSN functions, it remains unclear if iMSN may have specialized functions in inhibiting compulsive reward-seeking within each striatal subregion. Compulsive-like alcohol drinking refers to persistent alcohol drinking despite negative consequences, the hallmark of defining alcohol use disorder (AUD) in the Diagnostic and Statistical Manual of Mental Disorders 5 (DSM 5) (Grant et al., 2015). This compulsive aspect is a major obstacle in treating AUD (Dawson et al., 2005; Moos and Moos, 2006; Sinha, 2009; Koob and Volkow, 2010; Hopf and Lesscher, 2014). Thus, understanding how the activity of iMSN within each dorsal striatal region contributes to compulsive behavior may help guide the target and development of new therapeutic strategies.

Here, we selectively ablate iMSN circuits in the DMS and DLS to investigate their roles in compulsive reward-seeking and drinking behaviors using escalating concentrations of quinine adulterated ethanol solution in operant conditioning and two-bottle choice drinking tasks. Our findings indicate that the DLS iMSN function is essential in inhibiting compulsive-like behavior.

## Methods

### Animals

All experimental procedures were approved by the Mayo Clinic Institutional Animal Care and Use Committee and performed following National Institute of Health (NIH) guidelines. C57BL/6J male and female mice were purchased from Jackson Laboratory (Bar Harbor, ME). D1-tdT/D2-eGFP bitransgenic mice were bred in the lab. Mice were housed in standard Plexiglas cages. The colony room was maintained at a constant temperature (24 ± 1°C) and humidity (60 ± 2%) with a 12 hr light/dark cycle (lights on at 06:00 A.M.). We used 8- to 10-week-old mice for all experiments. Mice were allowed *ad libitum* access to food and water. For the operant conditioning tests, mice were food-restricted to 85% of their baseline weight, at which time they were maintained for the duration of experimental procedures.

### Operant conditioning

We conducted operant conditioning using the same operant chambers/schedules as our previous studies (Hong et al., 2019; Kang et al., 2020; Baker et al., 2023) (Supplemental Fig. S1B). Briefly, mice were placed in operant chambers (Med-Associates, St Albans, VT) in which they poked a single hole for an outcome of ethanol-containing solution (10 μl per reinforcement). For all experiments, sweetened ethanol (10% sucrose+10% ethanol, 10S10E) was used as the reward. Before training, mice were food-restricted to approximately 85% body weight, which was maintained for the duration of experimental procedures. On the first day, mice were trained to approach the reward magazine with a reward delivered on a random time schedule for 30 min. Next, mice were trained on a fixed ratio 1 schedule for 30 min in 7 sessions. After acquiring nose-poking behavior, mice were trained on random interval (RI10 1-2days/RI30 5 days/RI60 3 days) to develop habitual behavior or random ratio (RR2 1–2 days/RR5 5 days/RR10 3 days) to form goal-directed reward-seeking. Sessions were completed after 30 min or following 60 reward reinforcements. RI10/30/60 delivered one reward outcome on average every 10/30/60s after the last reward outcome to develop habitual reward-seeking. RR2/5/10 delivered one reward outcome on average every 2/5/10 response in the correct nose-poke hole to develop goal-directed reward-seeking.

### Reward devaluation and extinction test

On the devalued day, mice were given 1 hour of *ad libitum* access to the outcome (10S10E) previously earned by nose poking (for devaluation) or food pellets and then underwent serial nonreinforced extinction sessions in each training context. The order of the valuation context was counterbalanced across mice and was 10 min in duration. According to the sensory-specific satiety theory, we conducted reward devaluation for 1 hr and the extinction test for 10 min to determine which is more dominant between goal-directed behavior and habit. We provided unlimited food chow to make the reward worthwhile for the valued state, whereas we provided an unlimited 10S10E solution to devaluate the outcome value. We conducted the extinction tests after the last sessions of RR10, and RI60.

### Quinine adulteration test

Mice were placed in the RI60 operant conditioning task with increasing concentrations of quinine in the 10S10E solution (0, 100, 250 µM quinine). Mice performed two RI60 sessions at each quinine concentration and data was averaged between the two sessions.

### Stereotaxic surgery for virus injection

Mice were anesthetized with isoflurane (3.5% in oxygen gas) using a VetFlo^TM^ vaporizer with a single-channel anesthesia stand (Kent Scientific Corporation, Torrington, CT) and placed on the digital stereotaxic alignment system (model 1900; David Kopf instruments). Hair was trimmed, and the skull was exposed using 8-gauge electrosurgical skin cutter (KLS martin, Jacksonville, FL). The skull was leveled using a dual-tilt measurement tool. Holes were drilled in the skull at the appropriate stereotaxic coordinates. Viruses were infused to the DLS (AP + 0.7 mm, ML +/- 2.5 mm, DV –3.1 mm from bregma), DMS (AP + 0.7 mm, ML +/- 1.25 mm, DV –3.1 mm from bregma) or GPe (AP –0.4 mm, ML +/- 2.0 mm, DV −3.8 mm from bregma) at 100 nl/min for 4 minutes (400 nl total) through a 33-gauge injection needle (cat # NF33BV; World Precision Instruments) using a microsyringe pump (Model UMP3; World Precision Instruments). The injection needle remained in place for an additional 5 min following the end of the injection. All viruses were purchased from Addgene and were injected at the following titers: retrograde AAV-Ef1a-mCherry-IRES-Cre (7×10^12^ vg/mL), AAV5-flex-taCasp3-TEVp (7×10^12^ vg/mL), AAV-Ef1a-DIO-EYFP (7×10^12^ vg/mL). Following stereotaxic surgery, we injected buprenorphine extended release (1 mg/kg, S.C.; ZooPharm, Laramie, WY, USA) to alleviate post-surgery pain.

### Caspase 3-mediated circuit ablation

To ablate the circuit in a targeted fashion, we used a genetically engineered caspase 3 (Casp3), whose activation commits a cell to apoptosis (Chelur and Chalfie, 2007). Endogenous caspase 3 exists as procaspase 3, which is cleaved into its active form by upstream apoptotic signals and other caspase proteins. This genetically engineered caspase lacks the cleavage site for upstream caspases and can only be cleaved by a tobacco etch virus protease (TEVp) which is coexpressed in an AAV (Yang et al., 2013). AAV-flex-taCasp3-TEVp was expressed in a cre-dependent manner in the DLS to only ablate cells that express Cre recombinase from retrograde injections in the GPe (AAV-Ef1a-mCherry-IRES-Cre). Importantly, caspase 3 triggers cell-autonomous apoptosis, minimizing the risk of off-target effects and toxicity to neighboring cells (Mallet et al., 2002; Smart et al., 2017; Patton et al., 2021).

### Immunofluorescence

Brains were fixed with 4% paraformaldehyde (Sigma-Aldrich, St. Louis, MO) and transferred to 30% sucrose (Sigma-Aldrich) in phosphate-buffered saline at 4 °C for 72 hr. Brains were then frozen in dry ice and sectioned at 40 µm using a microtome (Leica Corp., Bannockburn, IL). Brain slices were stored at −20 °C in a cryoprotectant solution containing 30% sucrose (Sigma-Aldrich) and 30% ethylene glycol (Sigma-Aldrich) in phosphate-buffered saline. Sections were incubated in 0.2% Triton X-100 (Sigma-Aldrich), 5% bovine serum albumin in phosphate-buffered saline for 1 hr, followed by incubation with the primary antibody in 5% bovine serum albumin overnight at 4 °C. After three washes in phosphate-buffered saline, the sections were mounted onto a glass slide coated with gelatin and cover-slipped with a VECTASHIELD® antifade mounting medium (Vector Laboratories, Burlingame, CA). Images were obtained using an LSM 780 laser scanning confocal microscope (Carl Zeiss, Heidelberg, Germany) using a 10x or 40x water-immersion lens. Cell count values were determined using Image J (1.53t, National Institute of Health, USA). We used 509 (eGFP), 581 (tdTomato), 610 (mCherry), and 405 channels for imaging.

### Open field test

The open-field test (OFT) was conducted in chambers (Med-Associates, St Albans, VT) to measure the locomotor responses of mice. The session lasted 30 minutes and total distance and velocities were recorded using beam breaks. Animals were habituated to the testing room in their home cage 1 hour prior to behavioral testing.

### Two-bottle choice continuous access ethanol-drinking

Oral ethanol-containing solution (10S10E) consumption and preference were examined using a two-bottle choice test in the mouse home cage. Mice were individually housed and given 24 hr access to two bottles: water and 10S10E. Every other day, the bottles were measured, and bottle positions were switched to prevent place preference development. After baseline consumption and preference were established for 8 days, mice underwent two surgeries 1 week apart. After 3 weeks of the second injection, mice were single-housed in the home cage for the two-bottle choice test. Mice were given 24 hr access to two bottles: water and 10S10E but with increasing concentrations of quinine in the 10S10E solution (0, 100, 250 µM quinine). Mice performed two sessions (4 days, 2 days on each side) at each quinine concentration and data was averaged between the two sessions. 10S10E and water consumption were normalized for evaporation (Nam et al., 2013; Leon et al., 2023). Briefly, the total volume of the liquid evaporated was calculated by averaging 2-days of evaporation from the four control cages without mice. Then, the volume of water or ethanol that evaporated was subtracted from the total consumption of water or ethanol for each mouse.

### Data analysis

All data are represented as mean ± SEM using Prism 9.0 (GraphPad Software, San Diego, CA). The statistical significance was set at *p* < 0.05. The statistical significance was set at *p* < 0.05. All *p* values less than 0.05 were marked as “*” in the graphs. Detailed statistical tests and data with exact *p* values are listed in Supplemental Table S1.

## Results

### iMSNs primarily project to the GPe

The dorsal striatal projections can be divided into two distinct pathways: direct pathway medium spiny neurons (dMSN) target the entopeduncular nucleus (EP; rodent homolog of the primate internal globus pallidus), and substantia nigra pars reticulata (SNr) directly, whereas indirect pathway medium spiny neurons (iMSN) project to the external part of the globus pallidus (GPe), which then send axons to the output nuclei via the subthalamic nucleus (STN) (Alexander and Crutcher, 1990). To measure the effect of ablation, we injected retrograde mCherry virus into the GPe and counted the mCherry+ cells in the DLS and DMS of both control and casp3 ablated mice. We found that casp3 ablation indeed significantly reduces the neurons present within the DLS and DMS (Supplemental Fig. S2B and S2D, Unpaired *t* test, *p <* 0.05). Several studies have revealed that dMSN has axon collaterals in the GPe (Fujiyama et al., 2011; Kawaguchi et al., 1990; Wu et al., 2000). To this, we injected the retrograde mCherry virus into the GPe and counted iMSN in the DMS and DLS of both control and casp3 mice of bitransgenic D1-tdT/D2-eGFP mice. We found a significantly reduced number of iMSN in both DLS and DMS with a greater ablation of iMSN in the DLS (Fig. 1B and 1D, Unpaired *t* test, *p <* 0.05). This suggests that iMSN from the dorsal striatum primarily project to the GPe and the effect on behavior we observe is due to the partial ablation of these iMSN.

**Figure 1.**
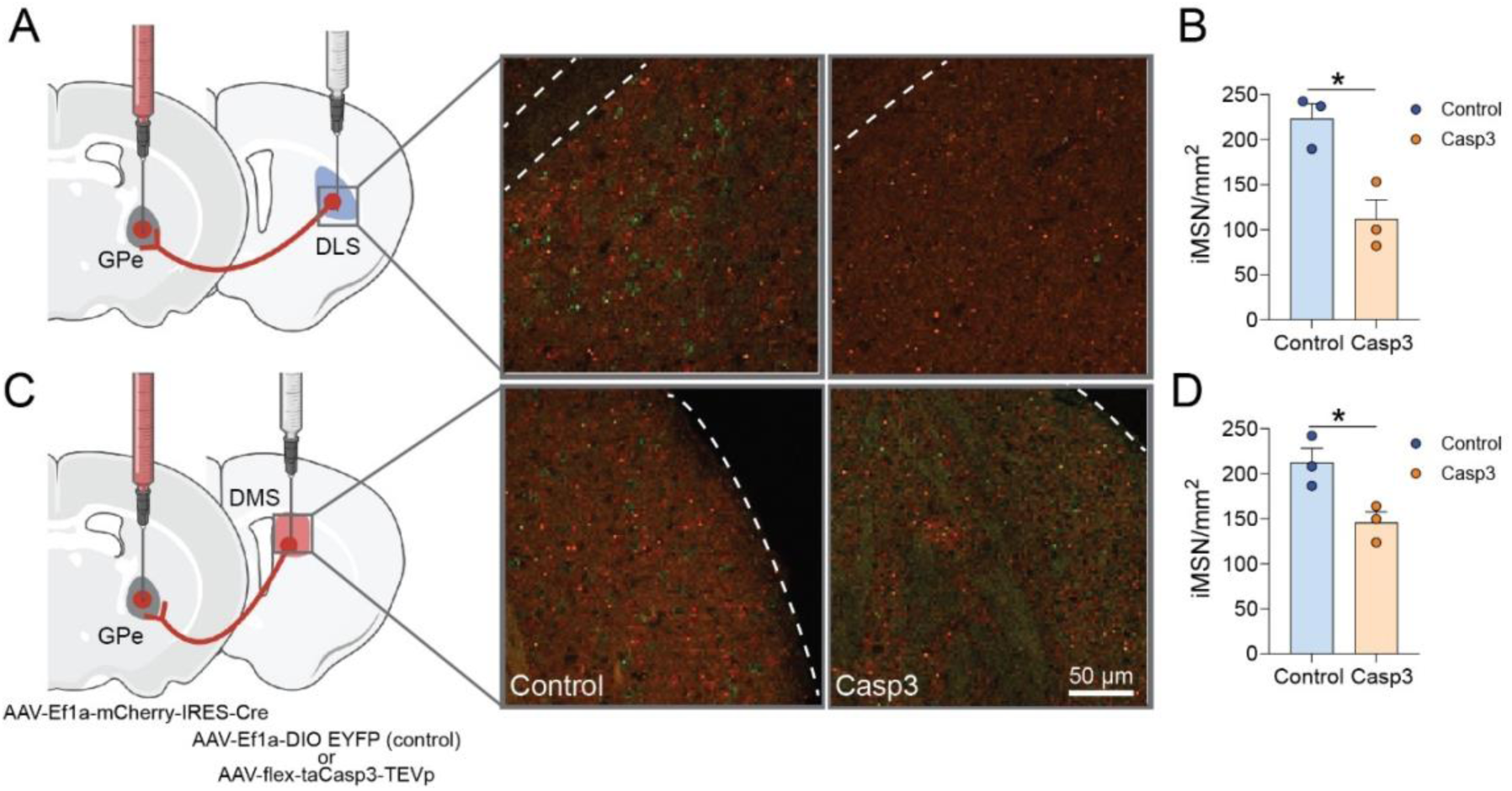
Effect of Caspase 3 (casp3)-dependent ablation of dorsal striatum iMSN. ***A***, Schematic of virus injections for Cre-dependent ablation of DLS neurons and representative DLS images, dMSN (red) iMSN (green) of sham control and caspase mice (Scale: 50 µm) from *n* = 3 mice/group. ***B***, Caspase mice showed a significant reduction of iMSN in the DLS (t = 3.87, *p* = 0.018). ***C***, Schematic of virus injections for Cre-dependent ablation of DMS neurons and representative DMS images, dMSN (red) iMSN (green), of sham control and caspase mice (Scale: 50 µm) from *n* = 3 mice/group. ***D***, Caspase mice showed a significant reduction of iMSN in the DMS (t = 3.32, *p* = 0.029). Figure (***A***) and (***C***) was created with BioRender.com.

### Caspase3-dependent ablation of ***iMSN^DLS→GPe^*** promotes increased compulsive-like ethanol-seeking but not motor suppression

To determine whether ablation of the iMSN^DLS→GPe^ circuit modulates goal-directed or habitual behavior, we used a Cre-dependent caspase 3 virus, which induces cell-autonomous death with minimal toxicity to neighboring cells (Yang; Mallet et al., 2002; Chelur and Chalfie, 2007; Smart et al., 2017; Patton et al., 2021). We bilaterally injected a mCherry-tagged retrograde virus expressing Cre recombinase [AAV-Ef1a-mCherry-IRES-Cre] into the GPe, followed by a second injection of Cre-dependent caspase-3 (AAV5-flex-taCasp3-TEVp; or AAV5-Ef1a-DIO EYFP (control) into the DLS (Supplemental Fig. S1A). We validated a significant reduction in mCherry-positive neurons in the DLS of the caspase group (unpaired *t-*test, *p* < 0.05; Supplemental Fig. S2B).

Since the DLS has long been associated with motor function, we examined if DLS iMSN ablation resulted in motor dysfunction. In the open field test (Supplemental Fig. S3A), we observed no significant differences in spontaneous locomotion, velocity, or time spent in the center zone, indicating that DLS iMSN ablation does not alter basic motor function (Mann-Whitney *U* test, *p* > 0.05; Supplemental Fig. S3B).

In the devaluation test, both control and caspase mice in the RR group showed significant reduction in nose pokes for the devalued state (Wilcoxon test, *p* < 0.05; Fig. 2B), consistent with goal-directed behavior. In contrast, both control and caspase mice in the RI group showed no significant differences between the valued and devalued states (Wilcoxon test, *p* > 0.05; Fig. 2C) indicating habitual behavior. Altogether our results demonstrate iMSN circuit ablation does not impair overall performance or acquisition rate in the operant reward-seeking task.

**Figure 2.**
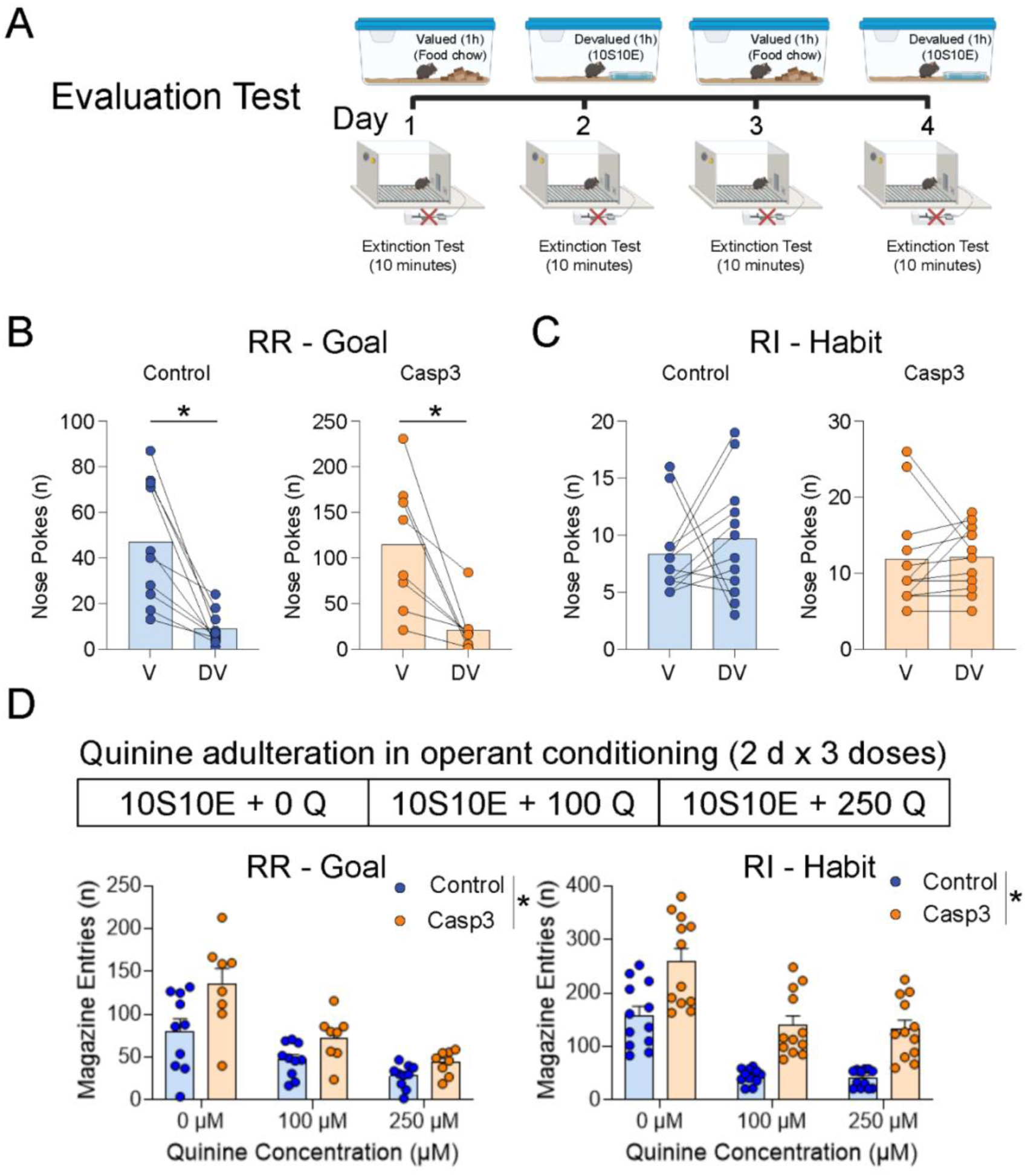
Effect of Caspase 3 (casp3)-dependent ablation of dorsolateral striatum (DLS) indirect medium spiny neurons (iMSN) projecting to external globus pallidus (GPe) on goal-directed and habitual alcohol-seeking. ***A***, Schematic of evaluation test to determine reward-outcome valuation testing. ***B***, Control and Casp3 animals showed significantly decreased nosepoke behavior in the devalued state, indicating goal-directed behavior. ***C***, Control and Casp3 animals showed no differences between the valued and devalued states, indicating habitual behavior. ***D***, Casp3 mice showed increased compulsive ethanol-seeking behaviors in quinine adulterated ethanol-seeking tasks. Wilcoxon test (**B-C**). *n* = 8 −12/group. Data represent mean ± SEM. Two-way repeated measures ANOVA with Tukey’s *posthoc* tests were used for (***D***). See Table S1 for full statistical information. Figure *(**A**)* was created with BioRender.com.

Next, we examined whether the iMSN^DLS→GPe^ ablation influenced compulsive-like seeking behavior. In the quinine adulteration test, all mice displayed reductions in reward-seeking behaviors as the quinine concentration increased (RM Two-way ANOVA, *p* < 0.05; Fig. 2D), showing sensitivity to the aversive taste of quinine. However, caspase mice showed increased magazine entries compared to control mice in both RR and RI groups, indicating more compulsive-like seeking behaviors (RM Two-way ANOVA, *p* < 0.05; Fig. 2D). Collectively, this indicates that decreased iMSN^DLS→GPe^ activity contributes to the resilience to bitter tasting quinine containing ethanol solution, suggestive of compulsive-like ethanol seeking behaviors.

### Caspase3-dependent ablation of iMSN^DMS→GPe^ shifts mice toward habitual-behavior

To determine whether ablation of iMSN^DMS→GPe^ modulates goal-directed or habitual reward-seeking, we injected a mCherry-tagged retrograde Cre-virus into the GPe, followed by a second injection of Cre-dependent caspase-3 (or control virus) into the DMS (Supplemental Fig. S2C). We saw a significant reduction in mCherry-positive neurons in the DMS in the caspase group (unpaired *t* test, *p* < 0.05; Supplemental Fig. S3D).

Previous studies have implicated iMSN in the DMS being associated with motor function (Kravitz et al., 2012; Cui et al., 2021), so we sought to determine if partial iMSN ablation resulted in motor dysfunction. In the open field test (Supplemental Fig. 3A), we observed no significant differences in spontaneous locomotion, velocity, or time spent in the center zone, indicating that DMS iMSN ablation does not alter basic motor function (Mann-Whitney test, *p* > 0.05; Supplemental Fig. S3C).

In the devaluation test, control mice in the RR group showed a reduction in extinction session nose pokes for the devalued state (Wilcoxon test, *p* < 0.05; Fig. 3B), consistent with goal-directed behavior. Consistent but conversely to our previous findings where activation of iMSN in DMS shifted habitual to goal-directed behavior (Kang; Kang et al., 2020), we find interestingly, RR caspase mice exhibit habitual behavior during devalued state (*p* > 0.05; Fig. 3B), indicating that loss of iMSN^DLS→GPe^ circuit function promotes a shift from goal-directed to habitual behavior. In contrast, RI caspase mice showed no significant differences between the valued and devalued states in both control and caspase mice (*p* > 0.05; Fig. 3C). Overall, this indicates that DMS iMSN ablation specifically reduces goal-directed behavior without changing habitual behaviors during conditioned reward.

**Figure 3.**
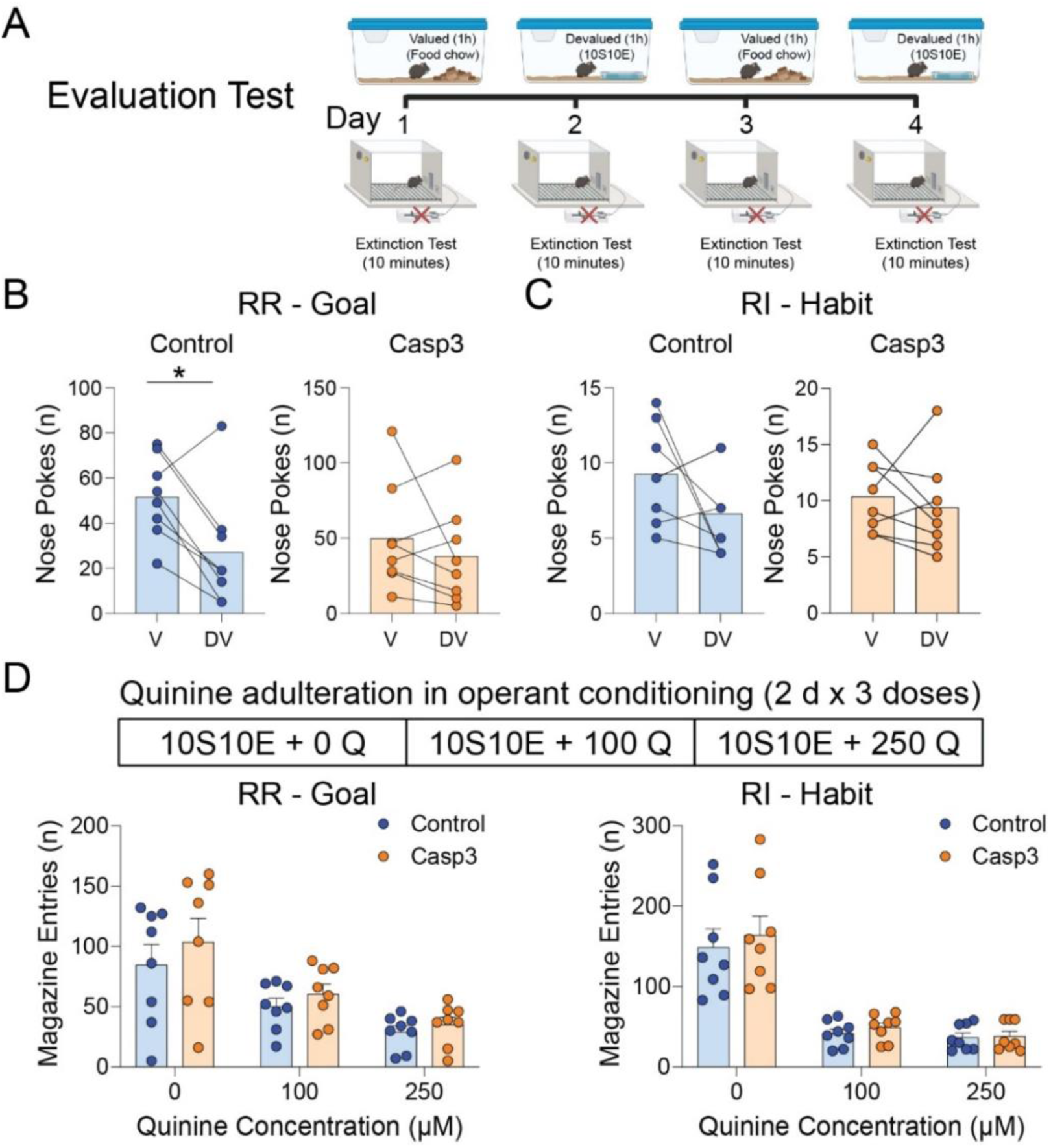
Effect of Caspase 3 (casp3)-dependent ablation of dorsolateral striatum (DLS) indirect medium spiny neurons (iMSN) projecting to external globus pallidus (GPe) on goal-directed and habitual alcohol-seeking. ***A***, Schematic of evaluation test to determine reward-outcome valuation testing. ***B***, Control mice showed significantly decreased nose poke behavior in the devalued state, indicating goal-directed behavior. However, Casp3 mice showed a shift to habitual behavior. ***C***, Control and Casp3 animals showed no differences between the valued and devalued states, indicating habitual behaviors. ***D***, Control and Casp3 mice showed no difference in compulsive ethanol-seeking behaviors in quinine adulterated ethanol-seeking tasks. Wilcoxon test (***B-C***). *n* = 8/group. Data represent mean ± SEM. Two-way repeated measures ANOVA with Tukey’s *posthoc* tests were used for (***D***). See Table S1 for full statistical information. Figure (***A***) was created with BioRender.com.

### Caspase3-dependent ablation of iMSN^DLS^**^→^**^GPe^ promotes compulsive ethanol-drinking in mice

Recurrent use of ethanol despite negative consequences is a hallmark feature of compulsive alcohol drinking in AUD. Given that we found increased compulsive-like alcohol-seeking in iMSN^DLS→GPe^ ablated mice, we asked whether iMSN ablation would increase voluntary ethanol intake and preference. We first established baseline ethanol intake which showed distinct sex-specific drinking behaviors. While both, male and female, mice show no difference in preference for ethanol, female mice exhibit higher consumption of ethanol (Supplemental Fig. S4). We then examined the effect of partial ablation of iMSN from the two dorsal striatal regions. To this, we subjected 10 mice (5 male + 5 female) per group to the two-bottle choice continuous access ethanol drinking test. We found that increasing concentrations of quinine reduced ethanol consumption and preference for both caspase and control groups showing sensitivity to bitter taste of quinine. Consistent with our previous findings, we found that iMSN^DLS→GPe^ ablated mice showed higher preference than the control group (RM Two-way ANOVA, Fig. 4B). While we did not see a group difference in consumption between the caspase and control group for the iMSN^DLS→GPe^ ablated mice, we did see a significantly higher consumption in the caspase group at 250 µM concentration of quinine, again showing resilience to the bitter tasting ethanol solution exhibiting more compulsive-like behavior. The iMSN^DMS→GPe^ control and caspase group showed no group differences in ethanol preference or consumption (RM Two-way ANOVA, Fig. 4C). Together, these results suggest that reduced iMSN^DLS→GPe^ activity contributes to compulsive-like ethanol drinking.

**Figure 4.**
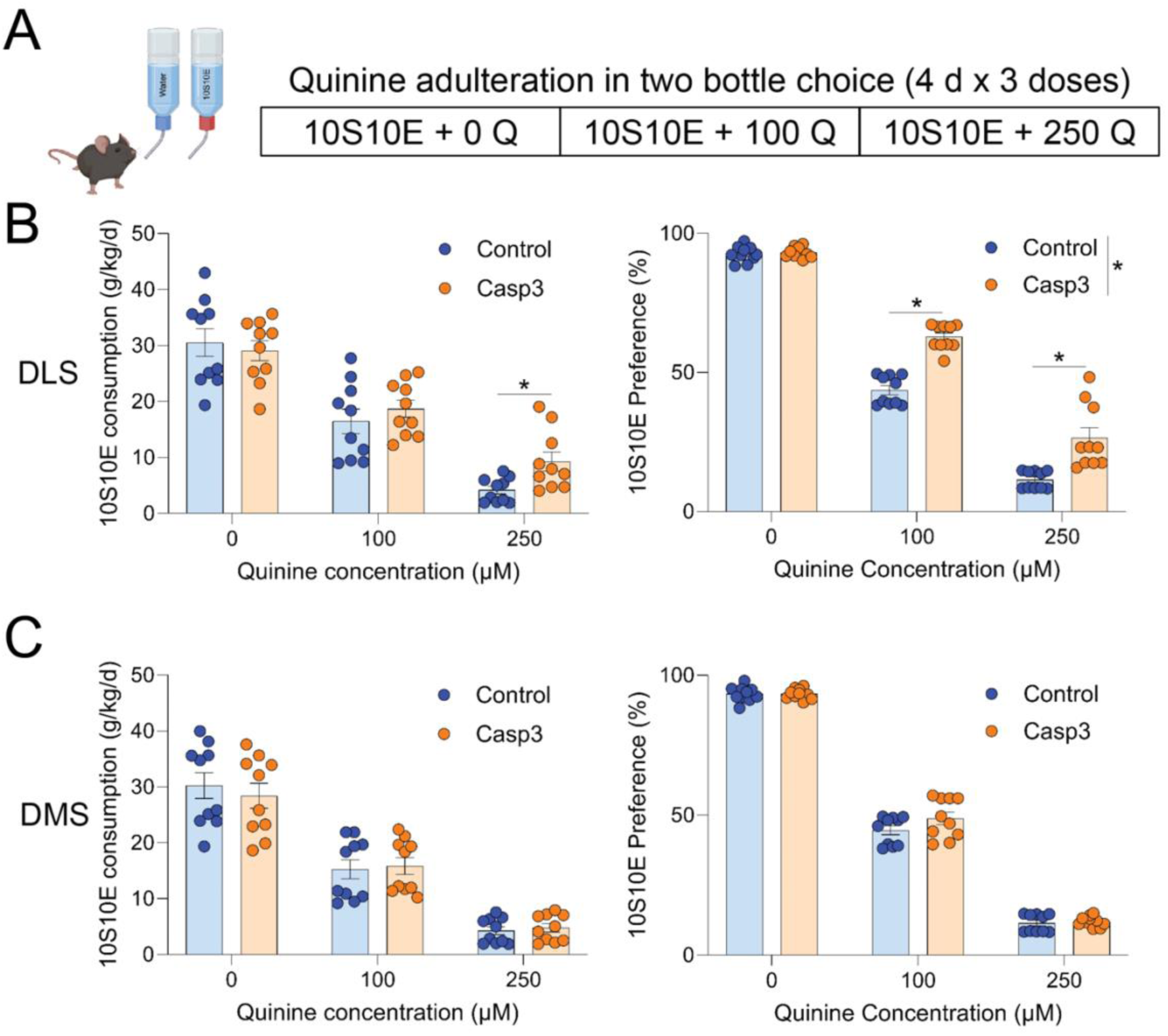
Effect of Caspase 3 (casp3)-dependent ablation of dorsal striatum indirect medium spiny neurons (iMSN) projecting to external globus pallidus (GPe) on ethanol-drinking. ***A***, Two-bottle choice continuous access paradigm showing the increasing concentration of quinine in 10S10E. ***B***, Partial ablation of iMSN^DLS→GPe^ (Casp3 mice) promotes compulsive ethanol-drinking behaviors in C57BL/6J mice. ***C,*** Partial ablation of iMSN^DMS→GPe^ (Casp3 mice) showed no change in ethanol-drinking behaviors in C57BL/6J mice. *n* = 8/group. Data represent mean ± SEM. Two-way repeated measures ANOVA with Tukey’s *posthoc* tests were used for (***B***, ***C***). See Table S1 for full statistical information. Figure (***A***) was created with BioRender.com.

## Discussion

In this study, we demonstrate that DLS iMSN plays a critical role in suppressing compulsive-like behaviors. Specifically, we observed that partial ablation of iMSN^DLS→GPe^ promotes compulsive ethanol-seeking and drinking behaviors in C57BL/6J mice.

Based on other studies (Lobo and Nestler, 2011; Kravitz et al., 2012), we hypothesized that DLS iMSN dysfunction would further promote reward-seeking behaviors. However, our results indicated that reward devaluation was not sensitive enough to show any changes in reward-seeking following circuit ablation. Previously, we showed that activation of astrocytes in the DMS increases iMSN activity, which leads to a shift from habitual to goal-directed behavior (Kang et al., 2020). Conversely, partial iMSN ablation in the DMS leads to a shift from goal to habit in mice trained in the random ratio (RR) paradigm (Figure 3B).

Previously, studies in humans have shown that hyperactive striatum is associated with compulsive behavior (Saxena and Rauch, 2000; Menzies et al., 2008). Similarly, preclinical studies using various mouse models have also revealed that a hyperactive striatum is involved in compulsive-like behavior (Welch et al., 2007; Ahmari et al., 2013; Burguiere et al., 2013). Hyperactive direct pathway or a reduction in activation of indirect pathway have long been suggested to be the underlying mechanisms of compulsive behaviors (Graybiel and Rauch, 2000; Saxena and Rauch, 2000; Maia et al., 2008). Additionally, an imbalance between the two pathways has been shown to be associated with compulsive drug-seeking (Lobo and Nestler, 2011; Bock et al., 2013; Roltsch Hellard et al., 2019). Consistent with previous studies, our study presents the notion that reduced indirect pathway activity increases compulsive-like seeking and drinking behaviors. The role of iMSN in the DLS are critical for regulating compulsive-like seeking and drinking behaviors. The ablation of these neurons disrupts the balance between the direct and indirect pathways, leading to the disinhibition of reward-driven actions. This imbalance manifests as increased compulsive-like behaviors, including excessive ethanol seeking and drinking.

Interestingly, we found a significant increase in magazine entries in the quinine adulteration task while there were no differences in acquisition of goal-directed or habitual behavior (Figure 2). Furthermore, DLS iMSN ablation led to disinhibition of the bitter reward and continued to show significantly higher ethanol consumption during the two-bottle choice continuous access (Figure 4). This suggests that segregating possible effects on reward-seeking and reward-consumption may be critical to understanding unique effects in different behavioral models of habitual or compulsive behavior (Pegg et al., 2021; Wright and Wesson, 2021). The ability to change behavior in response to signals from the environment and to stop previously effective actions is known as behavioral flexibility (Kim and Hikosaka, 2015; Uddin, 2021). It facilitates problem-solving and enables modification of behavior to change the environment. According to Griffin and Guez (Griffin and Guez, 2014), behavioral flexibility includes a variety of skills, such as the capacity to perform an existing behavior in a novel setting, block a previously rewarded activity, and create new behavior (Griffin and Guez, 2014; Kim and Hikosaka, 2015; Uddin, 2021). Since behavioral flexibility is the ability to stop or control unnecessary or risky actions, proper activation of iMSN is required for goal-directed behaviors and voluntary movements. Overall, our findings support striatal iMSN role in reward aversion and compulsive-like behavior.

In conclusion, our results suggest that iMSN inputs are a critical mechanism for controlling the expression of compulsive behaviors toward alcohol use. The partial ablation of iMSN in the dorsal striatum leads to increased compulsive-like seeking and drinking behavior by reducing inhibitory control, which in turn leads to reduced behavior flexibility.

## Conflict of Interest

All the authors declare that the research was conducted without any commercial or financial relationships that could be construed as a potential conflict of interest.

## Acknowledgment

We thank all the laboratory members for their helpful discussion and comments. Figures were created with BioRender.com. This research was supported by the Samuel C. Johnson for Genomics of Addiction Program at Mayo Clinic, the Ulm Foundation, and the National Institute of Health (AA029258, AG072898).

## Author Contributions

**Humza Haroon:** Conceptualization, Investigation, Formal analysis, Writing – Original Draft, Writing – Review & Editing, Visualization. **Matthew Baker:** Conceptualization, Writing – Review & Editing, Visualization. **Doo-Sup Choi:** Resources, Writing – Review & Editing, Supervision, Project administration, Funding acquisition.

## Notes

Conflict of Interest: The authors declare no competing financial interests.

### Competing Interest Statement

The authors have declared no competing interest.

