## Supplemental Materials for "Caspase-dependent ablation of indirect medium spiny neurons projecting to external globus pallidus promotes compulsive ethanol-seeking and drinking behaviors"

Conflict of Interest: The authors declare no competing financial interests.

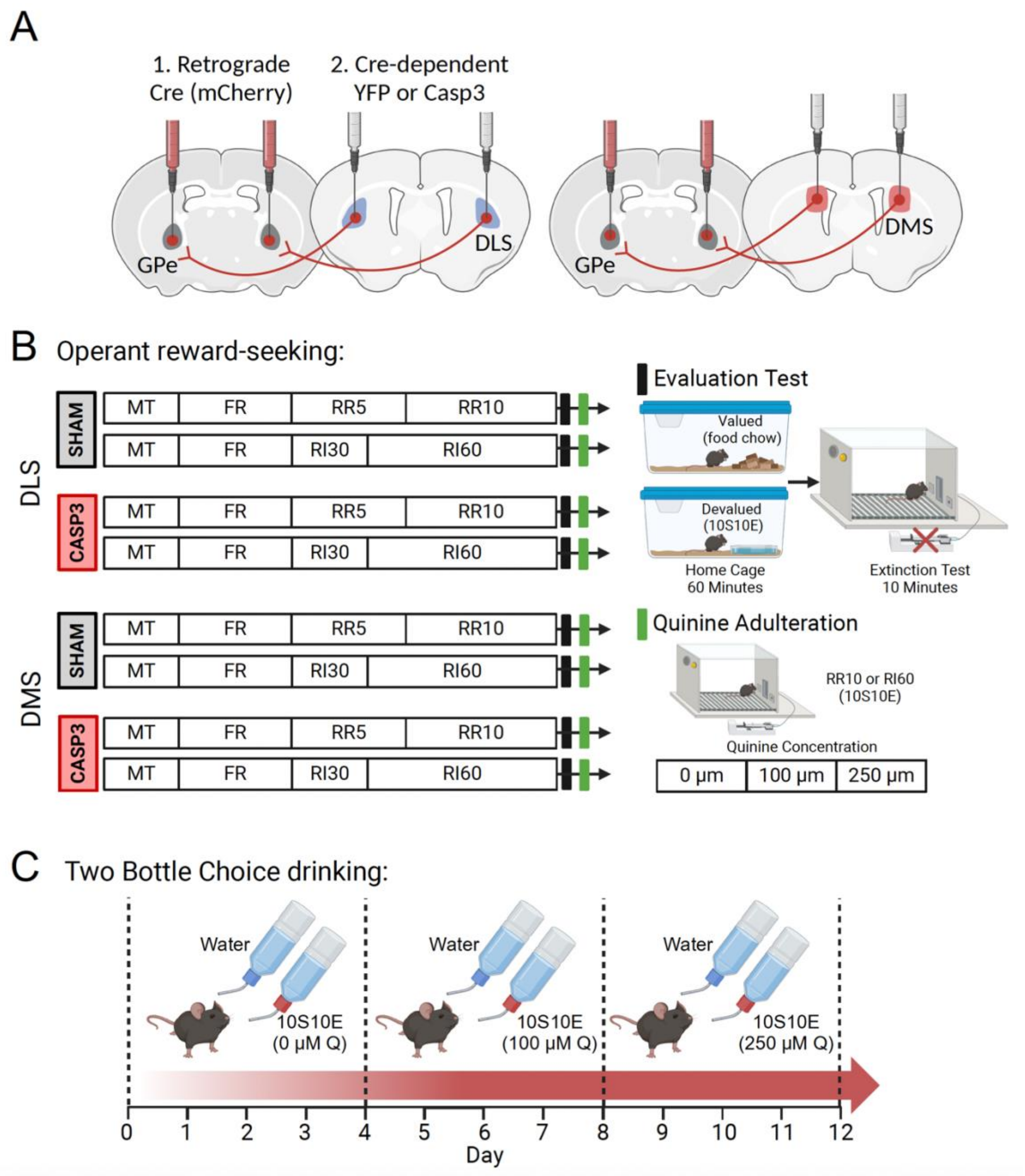


**Supplemental Figure S1.** Experimental Overview. ***A***, Schematic of retrograde cre- injection into the GPe, followed by cre-dependent Casp3 injection into the DLS. ***B****,* Operant behavior schedule to develop habitual seeking behavior. After the last day of testing devaluation testing was performed, followed by additional RI60 sessions with increasing quinine reward concentrations every 2 sessions for 6 days. ***C****,* Two-bottle choice continuous access paradigm with increasing quinine reward concentrations every 4 days. All figures were created with BioRender.com.

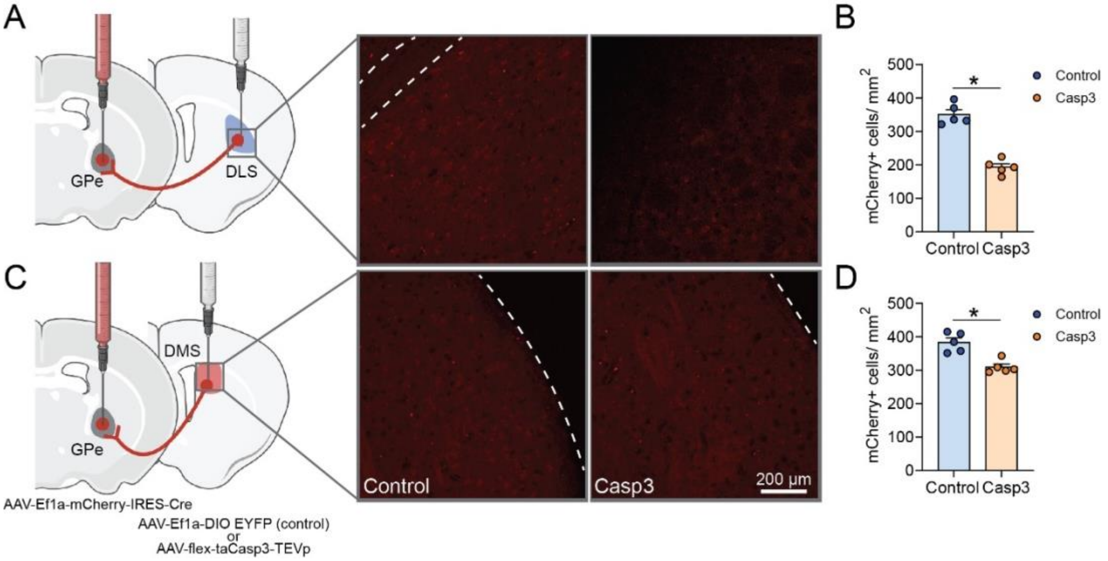


**Supplemental Figure S2**. Effect of Caspase 3 (casp3)-dependent ablation of dorsal striatum iMSN. ***A***, Schematic of virus injections for Cre-dependent ablation of DLS neurons and representative DLS images of sham control and caspase mice (Scale: 200 µm) from n = 5 mice/group. ***B***, Caspase mice showed a significant reduction of mCherry-positive neurons in the DLS ($t = 9.24$, $p < 0.0001$). ***C***, Schematic of virus injections for Cre-dependent ablation of DLS neurons and representative DMS images of sham control and caspase mice (Scale: 200 µm) from $n = 5$ mice/group. ***D***, Caspase mice showed a significant reduction of mCherry-positive neurons in the DMS ($t = 4.73$, $p = 0.0015$). Figure (***A***) and (***C***) was created with BioRender.com.

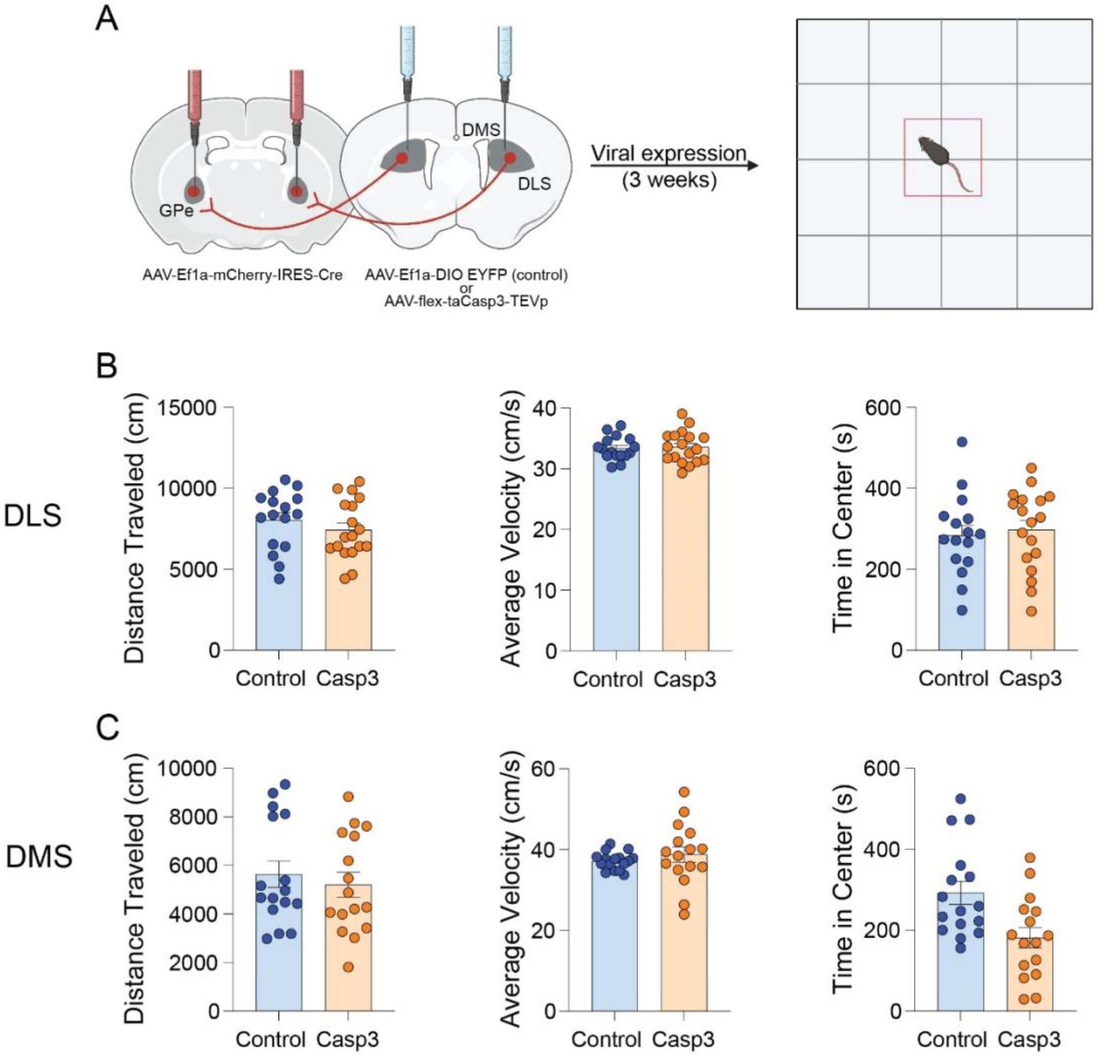


**Supplemental Figure S3.** Effects of Caspase 3 (casp3)-dependent ablation of iMSN on spontaneous locomotion in the open field test. ***A***, Schematic of virus injection followed by OFT. ***B***, DLS iMSN ablation, and ***C***, DMS iMSN ablation did not alter the distance traveled or average velocity or time in the center zone. Figure (***A***) was created with BioRender.com.

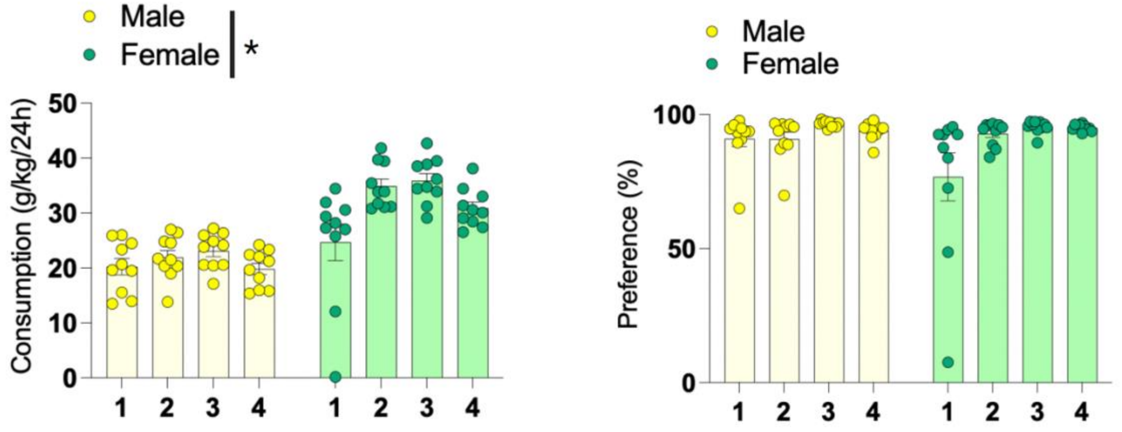


**Supplemental Figure S4**. Baseline drinking in male and female mice before virus injection. Female mice show higher consumption than male mice (left) with no difference in preference (right). *n* = 10/group. Data represent mean ± SEM. Two-way repeated measures ANOVA with Tukey's *posthoc* tests were used. See Table S1 for full statistical information.

**Table**

**Table S1. Summary of statistical analyses and results.**

| Figure | Statistical Tests | Comparison | Value | *p* value |
|---|---|---|---|---|
| Figure 1b | Unpaired *t*-test | Control vs Casp3 | t(4) = 3.865 | *p*=0.0181 |
| Figure 1d | Unpaired *t*-test | Control vs Casp3 | t(4) = 3.319 | *p*=0.0294 |
| Figure 2b | Wilcoxon test | RR Control: V vs DV | W = -55.00 | *p* = 0.002 |
| | | RR Casp3: V vs DV | W = -36.00 | *p*=0.0078 |
| Figure 2c | Wilcoxon test | RI Control: V vs DV | W = -26.00 | *p*=0.3398 |
| | | RI Casp3: V vs DV | W = 6.000 | *p*=0.7109 |
| Figure 2d | Two-way RM ANOVA | RR-ME: Conc | $F_{2, 18}$ = 64.23 | *p*<0.0001 |
| | | RR-ME: Group | $F_{1, 16}$ = 5.875 | *p*=0.0276 |
| | | RR-ME: Interaction | $F_{2, 32}$ = 4.890 | *p*= 0.014 |
| | | Tukey's posthoc: 0 | q = 3.308 | *p*=0.0348 |
| | | Tukey's posthoc: 100 | q = 3.201 | *p*=0.0427 |
| | | Tukey's posthoc: 250 | q = 3.261 | *p*=0.0361 |
| | | RI-ME: Conc | $F_{1, 23}$ = 104.4 | *p*<0.0001 |
| | | RI-ME: Group | $F_{1, 22}$ = 25.09 | *p*<0.0001 |
| | | RI-ME: Interaction | $F_{2, 44}$ = 0.093 | *p*=0.9112 |
| Figure 3b | Wilcoxon test | RR Control: V vs DV | W = -29.00 | *p*=0.0469 |
| | | RR Casp3: V vs DV | W = -6.000 | *p*=0.7422 |
| Figure 3c | Wilcoxon test | RI Control: V vs DV | W = -26.00 | *p*=0.0859 |
| | | RI Casp3: V vs DV | W = -12.00 | *p=0.4375* |
| Figure 3d | Two-way RM ANOVA | RR-ME: Conc | $F_{1, 16}$ = 33.82 | *p*<0.0001 |
| | | RR-ME: Group | $F_{1, 14}$ = 1.30 | *p*=0.4219 |
| | | RR-ME: Interaction | $F_{2, 28}$= 0.3632 | *p*=0.6987 |
| | | RI-ME: Conc | $F_{1, 14}$ = 80.35 | *p*<0.0001 |
| | | RI-ME: Group | $F_{1, 14}$= 0.2558 | *p*=0.6209 |
| | | RI-ME: Interaction | $F_{2, 28}$= 0.3632 | *p*=0.8145 |
| Figure 4b | Two-way RM ANOVA | Consumption: Conc | $F_{1, 26}$ = 1.83 | *p*<0.0001 |
| | | Consumption: Group | $F_{1, 10}$ = 1.30 | *p*=0.3953 |
| | | Consumption: Interaction | $F_{1, 18}$ = 0.47 | *p*=0.0108 |
| | | Tukey's posthoc: 0 | q = 0.6789 | *p*=0.6376 |
| | | Tukey's posthoc: 100 | q = 1.194 | *p*=0.4107 |
| | | Tukey's posthoc: 250 | q = 3.991 | *p*=0.0154 |
| | Two-way RM ANOVA | Preference: Conc | $F_{1, 26}$ =1077 | *p*<0.0001 |
| | | Preference: Group | $F_{1, 18}$ =39.78 | *p*<0.0001 |

| | | | | |
|---|---|---|---|---|
| | | Preference: Interaction | $F_{2, 36}$ =18.81 | *p*<0.0001 |
| | | Tukey's posthoc: 0 | q = 0.7083 | *p*=0.6234 |
| | | Tukey's posthoc: 100 | q = 12.43 | *p*<0.0001 |
| | | Tukey's posthoc: 250 | q = 5.595 | *p*=0.0025 |
| Figure 4c | Two-way RM ANOVA | Consumption: Conc | $F_{1, 24}$ = 374.1 | *p*<0.0001 |
| | | Consumption: Group | $F_{1, 18}$= 0.1422 | *p*=0.9057 |
| | | Consumption: Interaction | $F_{2, 36}$ =1.132 | *p*=0.3335 |
| | Two-way RM ANOVA | Preference: Conc | $F_{1, 19}$ = 2401 | *p*<0.0001 |
| | | Preference: Group | $F_{1, 18}$ = 1.865 | *p*=0.1888 |
| | | Preference: Interaction | $F_{2 , 36}$ = 1.888 | *p*=0.1660 |
| Figure S2b | Unpaired *t*-test | Control vs Casp3 | t(8) = 9.24 | *p*<0.0001 |
| Figure S2d | Unpaired *t*-test | Control vs Casp3 | t(8) = 4.728 | *p* = 0.0015 |
| Figure S3b | Mann-Whitney test | Distance: Control vs Casp3 | U = 22 | *p*= 0.54 |
| | | Velocity: Control vs Casp3 | U = 24 | *p*= 0.69 |
| | | Time: Control vs Casp3 | U = 70 | *p* = 0.23 |
| Figure S3c | Mann-Whitney test | Distance: Control vs Casp3 | U = 19 | *p*=0.1949 |
| | | Velocity: Control vs Casp3 | U = 29 | *p*=0.7984 |
| | | Time: Control vs Casp3 | U = 80 | *p* = 0.42 |
| Figure S4 | Two-way RM ANOVA | Consumption: Conc | $F_{1, 24}$ = 10.11 | *p*=0.0018 |
| | | Consumption: Group | $F_{1, 18}$= 0.1422 | *p*<0.0001 |
| | | Consumption: Interaction | $F_{3, 54}$ =4.015 | *p*=0.0119 |
| | | Tukey's posthoc, day 1 | q = 1.725 | *p* =0.245 |
| | | Tukey's posthoc, day 2 | q = 10.19 | *p*<0.0001 |
| | | Tukey's posthoc, day 3 | q = 10.96 | *p*<0.0001 |
| | | Tukey's posthoc, day 4 | q = 10.26 | *p*<0.0001 |
| | | Preference: Conc | $F_{1, 21}$ = 5.285 | *p*=0.0259 |

| | | | | |
|---|---|---|---|---|
| | Two-way RM ANOVA | Preference: Group | $F_{1, 18}$ = 1.100 | *p*=0.3081 |
| | | Preference: Interaction | $F_{18, 54}$ = 1.498 | *p*=0.0625 |
